# Post-Transcriptional Size-Dependent Expression of the Fission Yeast Cdc13 Cyclin

**DOI:** 10.1101/2023.01.16.524304

**Authors:** Samirul Bashir, Xi-Ming Sun, Yixuan Zhao, Nuria G. Martínez-Illescas, Isabella M. Gallego Lopez, Lauren Guerrero Negrón, Makoto Ohiro, Daniel Keifenheim, Tatiana Karadimitriou, Mary Pickering, Samuel Marguerat, Nicholas Rhind

## Abstract

The major fission yeast cyclin, Cdc13, has been shown to increase in concentration in correlation with cell size, and has been proposed to thereby regulate cell size at division. However, the mechanism of its cell-size regulation has been unknown. Here, we show that Cdc13 is regulated post-transcriptionally. Its transcript is not expressed in a size-dependent manner, rather a size-dependent concentration of protein is expressed from a size-independent concentration of mRNA. Moreover, we show that the expression of Cdc13 is, in fact, size dependent, as opposed to simply size-correlated due to time-dependent expression. We identify a 20-amino-acid motif, which includes the APC D-box degron, as necessary and sufficient for size-dependent expression, which allowed us to construct a size-independent allele of *cdc13*. Using this allele, we show that size-dependent expression of Cdc13 is not required for size control in fission yeast cells.

## Introduction

How a population of cells maintains a stable cell size is a fundamental question of cell biology for which no general answer has been discovered (Amodeo and Skotheim, 2016; Rhind, 2021). One potential solution to the problem of cell-size control is size-dependent expression of cell-cycle regulators (Fantes et al., 1975). Such proteins fall into two classes: diluted inhibitors and accumulating activators. Diluted inhibitors have been proposed to regulate size at the G1/S transition in budding yeast (Schmoller et al., 2015), algae (Liu et al., 2023), plants (D’Ario et al., 2021), and mammals (Zatulovskiy et al., 2020). In each case, an inhibitor of the G1/S transition—Whi5, TNY1, KRP4 and Rb, respectively—is proposed to be expressed at fixed number of molecules early in the cell cycle so that its concentration is dependent on the size of the cell, with small cells having a higher concentration of the inhibitor than large cells. The inhibitor has a concentration threshold above which it prevents the G1/S transition, so smaller cells must grow more in G1 to reach that threshold, but all cells will enter S at about the same size.

The other class of size-dependent regulators are accumulating activators, which are proposed to increase in concentration in proportion to cell size, such that they are only expressed at a sufficient concentration to drive cell cycle transitions when cells reach a critical size threshold. The existence of accumulating activators as regulators of size control was inferred from classic cell biology experiments (Prescott, 1956; Thormar, 1959; Herring, 1974; Fantes et al., 1975; Rhind, 2018). In the fission yeast, *Schizosaccharomyces pombe*, two mitotic activators—Cdc13, the B-type cyclin required for mitotic cyclin-dependent kinase (CDK) activity, and Cdc25, the tyrosine phosphatase required to activate CDK at the G2/M transition—have been shown to increase in concentration in G2 in correlation with cell size and have thus been proposed to be accumulating activators that regulate the G2/M transition (Moreno et al., 1990; Creanor and Mitchison, 1996; Keifenheim et al., 2017; Patterson et al., 2019; Curran et al., 2022; Miller et al., 2022; Miller et al., 2022).

Although size control by size-dependent expression of cell cycle regulators is an attractive paradigm, no such system has been shown to be required for cellular size control in any organism. In no system, and, in particular, in none of the systems described above, has abolition of size-dependent expression of a putative size-control protein led to loss of cellular size homeostasis. Therefore, at the very least, the proposed size-control mechanisms are redundant with other, as yet undiscovered, systems. Moreover, size-dependent expression is unusual—most proteins are expressed at a constant concentration as cells grow (Schmoller and Skotheim, 2015)—and in only one case, KRP4 in *Arabidopsis*, has a mechanism for size-dependent concentration been elucidated (D’Ario et al., 2021). Therefore, many important questions remain regarding if and how size-dependent protein expression regulates cell size.

We have addressed such questions in fission yeast, with particular focus on the major B-type cycle, Cdc13. We have investigated three major questions: Is Cdc13 expressed in a size-dependent manner? If so, how is Cdc13’s size-dependent expression regulated? and Is Cdc13’s size-dependent expression required for size control?

The first question is important because, in growing cells, size is correlated with time. So, a protein that increases in concentration as a cell grows larger could be directly regulated by cell size, or it could increase in concentration as a function of time after induction of expression, and only be indirectly correlated with size. One way to distinguish between size- and time-dependent expression of a protein is to uncouple the direct correlation between size and time in asynchronous cultures. In unperturbed cultures, there is a tight correlation between a cell’s size and the time it has grown since division. This correlation can be uncoupled by manipulating the size at division, so that cells in different cultures begin the cell cycle at different lengths (Miller et al., 2022). If one culture of cells divides at twice the size of another, a protein expressed at a size-dependent manner would be expected to be expressed at twice the concentration at the beginning of the subsequent cell cycle, whereas a protein expressed in a time-dependent manner would be expected to start at the same concentration in both cultures.

The second question we addressed is important because little is known about how proteins can be expressed in a size-dependent manner. Cdc25 size dependence is regulated at the transcriptional level (Keifenheim et al., 2017), although how its promoter is regulated by size is unknown. Cdc13 accumulates during S, G2 and metaphase (Creanor and Mitchison, 1996; Patterson et al., 2019; Miller et al., 2022) and then is degraded during anaphase and G1 by the anaphase-promoting complex (APC)-dependent proteolysis (Blanco et al., 2000). The APC recognizes Cdc13 via a short D-box motif in its N-terminus, although not in G2. Pcr1 and Atf1 transcriptionally regulate Cdc13 expression during S and G2 phases (Bandyopadhyay et al., 2017), but there is no indication that this regulation is size -specific.

The third question is important because in no case is the mechanism of cell size control understood. Cell-size control is an essential cellular function. Cells unable to coordinate growth and division are inviable, either because they divide too often and come inviably small or because they do not divide often enough and become inviably large. Therefore, abrogating cell size control is expected to lead to inviability. Loss of Rb leads to a loss of size dependence of the G1/S transition in mammalian cells (Zatulovskiy et al., 2020) and loss of Whi5 is presumed to have the same affect in budding yeast (Swaffer et al., 2021), but in neither case is cellular size control lost, presumably because of redundant size control mechanisms regulating other cell cycle transitions. A good candidate for such redundant regulation is size-dependent regulation of the G2/M transition. Therefore, it is important to understand how size affects this transition in general and how Cdc13 is involved in its regulation in fission yeast in particular.

## Results

### Cdc13 is Expressed in a Size-Dependent Manner

To investigate the level of Cdc13 expression in individual cells and to test for a correlation with cell size, we tagged the 3’ end of the endogenous *cdc13* gene with sequence encoding the green fluorescent protein NeonGreen (NG). Cdc13-NG localizes to the nucleus and is absent from binucleate post-anaphase cells, as expected (Booher et al., 1989), and cells expressing it show only a slight increase in length (12.8±2.9 μm for *cdc13-NG* cells (yFS1180) v. 11.8±2.5 μm for wild-type cells (yFS109)), demonstrating that the tagged protein is close to fully functional. We used widefield fluorescence microscopy to quantitate Cdc13-NG signal in cells and correlate it with cell length (Figure 1A).

**Figure 1.**
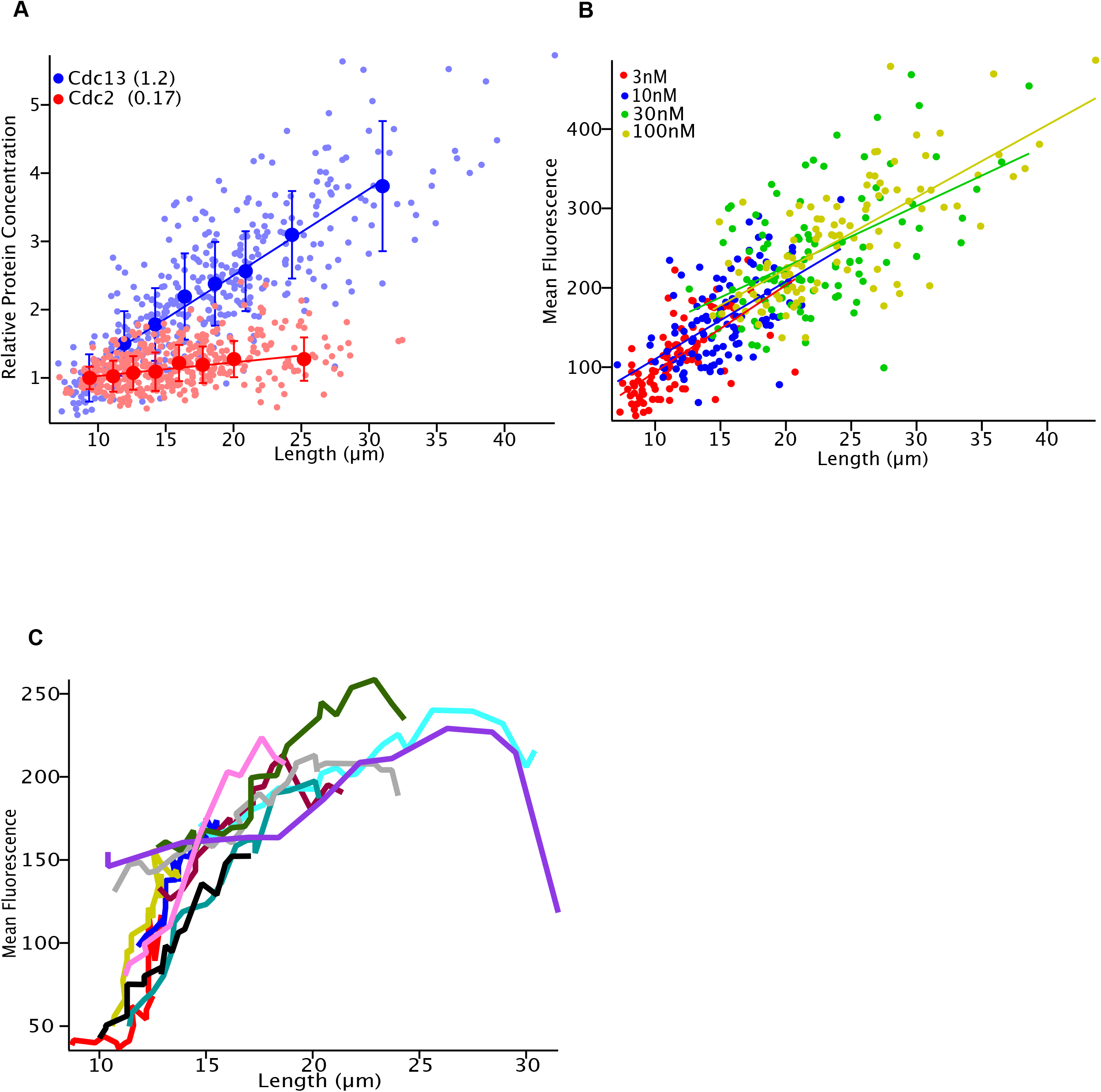
Cdc13 is Expressed in a Size-Dependent Manner. A) Cdc13 Concentration Correlates with Size. Asynchronous cells expressing Cdc13-NG from its endogenous locus (yFS1180) were analyzed by widefield fluorescence microscopy. Asynchronous cultures of cells spanning various size ranges were obtained by growth in 3nM, 10nM, 30nM, and 100nM estradiol to regulate the activity of Wee1 expressed from the *ZEV* promoter. Data from such cultures was combined and data from binucleate cells—which are in late mitosis, G1 and early S phase, and in which Cdc13 is degraded by APC-dependent proteolysis—was removed. Cdc2-GFP cells (yFS1133) were similarly analyzed. The small symbols represent individual cell; the large symbols represent an average of the data in 50-cell bins. The legend includes the size-correlation metric. B) Cdc13 is Expressed in a Size-Dependent Manner. Data from A was analyzed for individual cultures grown in different concentrations of estradiol and plotted in different colors. C) Time-lapse Analysis of Cdc13 Expression. Cdc13-NG (yFS1134) was imaged for 4 hours, and the mean nuclear intensity from 10 cells of different sizes was plotted.

We increased the range of cell length over which we measured Cdc13 concentration by overexpressing Wee1 using the ZEV estradiol-sensitive promoter, which allows us to control cell length with estradiol (Ohira et al., 2017). Wee1 is a kinase that inhibits CDK during G2. A high level of Wee1 prevents dephosphorylation of Cdc2 and arrests cell in G2 (Russell and Nurse, 1987). By varying the concentration of estradiol, we can titrate the specific activity of Wee1 to levels that result in asynchronous cultures with cell lengths that range from about 8 μm to 16 μm at 3 nM estradiol to those that range from about 15 μm to 30 μm at 100 nM estradiol (Figure 1A). These cultures are otherwise healthy and have doubling times between 2.5 hours and 3 hours, similar to wild-type cells (Russell and Nurse, 1987). By combining data from this range of sizes, we observed that Cdc13 concentration in cells correlates linearly with cell size over at least a 3.5-fold range (Figure 1A). We have also quantified Cdc13 in a strain in which the endogenous Cdc13 protein is tagged with an internal superfold GFP tag (Cdc13-sfGFP) (Chethan et al., 2025). Using this independent tag, we observed a similar expression pattern for Cdc13 (Figure S1A). In contrast, Cdc2 maintains a constant concentration over a similar size range (Figure 1A), as has been reported for almost every other protein in fission yeast (Marguerat et al., 2012; Curran et al., 2022).

As discussed in the introduction, the correlation between Cdc13 concentration and cell size could be directly due to size-dependent expression of Cdc13 or indirectly due to Cdc13 accumulating over time during the cell cycle. One way to distinguish between size- and time-dependent expression of Cdc13 is to uncouple the direct correlation between size and time in asynchronous cultures. We did so using the *ZEV:wee1* approach described above. Using four different concentrations of estradiol (3nM, 10nM, 30nM and 100nM), we produced four cultures that begin the cell cycle at a range of lengths (about 8 μm, 10 μm, 13 μm and 15 μm, respectively), but that all grow with the similar doubling times. By measuring Cdc13 concentration in these cultures, we find that Cdc13 correlates with the size of a cell, not the amount of time it has been growing since the beginning of the cell cycle. (Figure 1B). For instance, cells that are 15 μm long have about the same concentration of Cdc13 regardless of whether they are at the beginning of their cell cycle (as is the case for the culture grown in 30nM estradiol), the middle of their cell cycle (as is the case for the culture grown in 10nM), or at the end of their cell cycle (as is the case for the culture grown in 3nM). We further tested this hypothesis by performing time-lapse microscopy of Cdc13-NG in cells of different sizes. We analysed 10 cells spanning a range of sizes and imaged them for approximately 4 hours. Here, again, we observed that Cdc13 accumulation correlates with cell size rather than with time (Figure 1C). These results lead us to conclude that Cdc13 is expressed in a size-dependent manner.

One plausible mechanism for size-dependent expression of Cdc13 is that it is regulated by a positive feedback loop in which CDK activity increases the rate of Cdc13 synthesis. To examine this possibility, we expressed Cdc13-NG in combination with Cdc2-M17as, an allele that allows inhibition of Cdc2 kinase activity using the ATP analog 1NM-PP1 (Dischinger et al., 2008; Aoi et al., 2014). We found that disrupting the putative Cdc2-dependent positive feedback loop did not affect the size-dependent expression of Cdc13 (Figure S1B).

### Cdc13 Size-Dependent Expression is Regulated Post-Transcriptionally

To determine how Cdc13 size-dependent expression is achieved, we examined the size dependence of its transcript by measuring steady-state transcript levels in individual cells using single-molecule RNA fluorescence in situ hybridization (smFISH) (Keifenheim et al., 2017; Sun et al., 2020). As is the general case for transcripts in fission yeast and other organisms (Zhurinsky et al., 2010; Marguerat et al., 2012; Padovan-Merhar et al., 2015), the *cdc13* transcript maintains a constant concentration with respect to cell size (Figure 2). This result is observed in both in unperturbed, wild-type, asynchronous cells (Figure 2A) and in *ZEV:wee1* cells that vary over 5-fold in length from 4.1 to 28.8 μm in length. These results demonstrate that the size dependence of Cdc13 expression is achieved post-transcriptionally.

**Figure 2.**
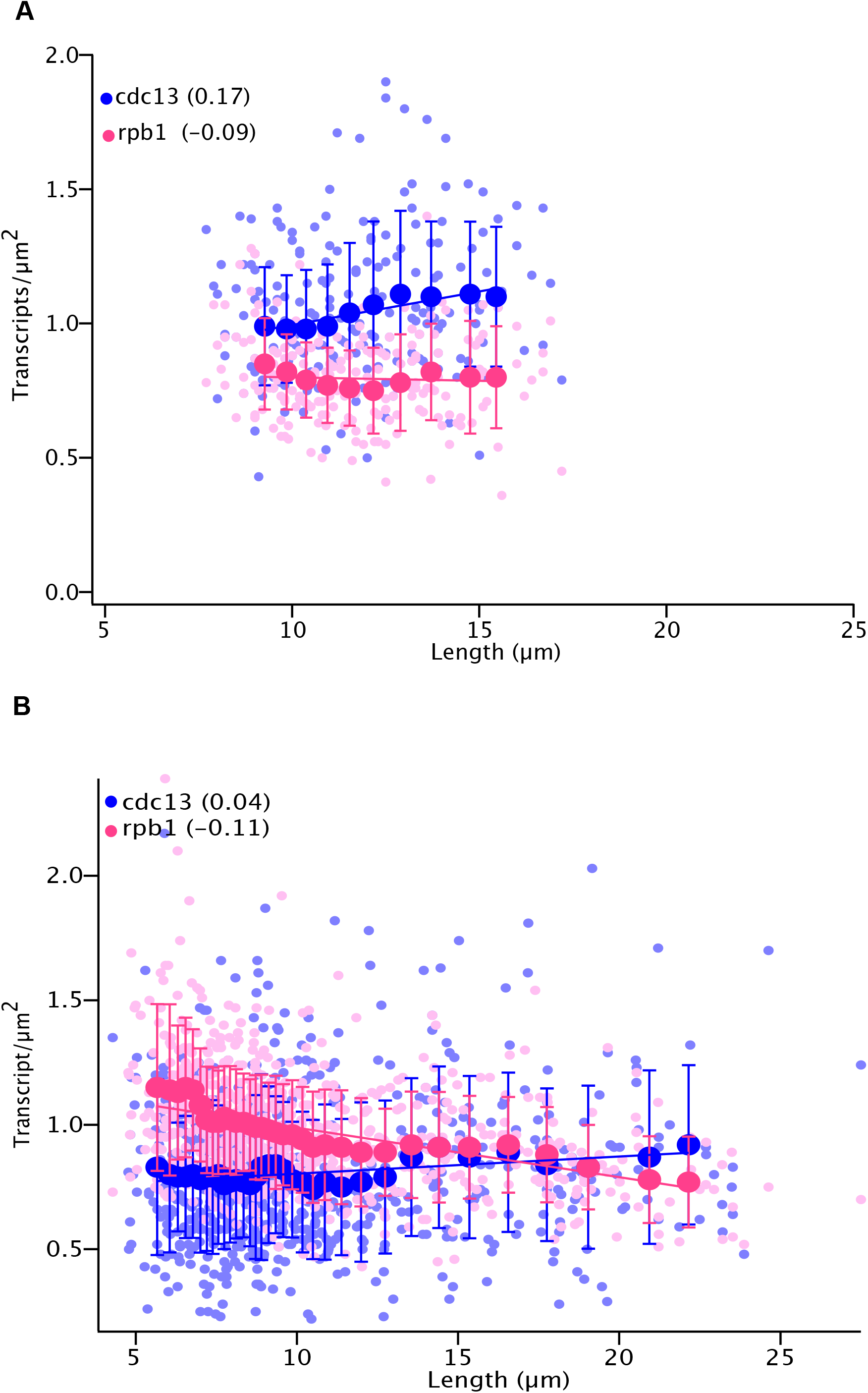
*cdc13* is not Transcribed in a Size-Dependent Manner. A) The *cdc13* Transcript Maintains a Constant Concentration in Asynchronous Wild-Type Cells. Wild-type cells (yFS105) were analyzed by smFISH for transcript numbers of *cdc13* and *rpb1*, a standard smFISH control. The small symbols represent individual cell; the large symbols represent the average of the data in 20-cell bins. The legend includes the size-correlation metric. B) The *cdc13* Transcript Maintains a Constant Concentration Over a Wide Range of Cell Sizes. Asynchronous cultures of *ZEV:wee1* cells (yFS970) spanning various size ranges were obtained by growth 0 to 100 nM beta-estradiol to regulate the expression of Wee1. *cdc13* and *rpb1* transcript numbers in individual cells were measured by smFISH. Data from all of the cultures was combined and presented as in A.

### Size-Dependence is Encoded in an N-Terminal Motif of the Cdc13 Protein

We next undertook a structure-function analysis of the *cdc13* gene to identify the region required for its size-dependent expression. Initially, we tested whether size dependence is encoded in the untranscribed, untranslated or translated parts of the gene. To do so in a way that allowed us to test essential parts of *cdc13*, we built an exogenous, NeonGreen-tagged copy of *cdc13* and integrated it into a strain expressing *ZEV:wee1*. Cdc13-NG expressed from its own promoter, with its own 5’- and 3’-UTRs, shows size dependence similar to expression from its endogenous locus (Figure 3A). Replacement of the *cdc13* promoter, 5’- and/or 3’-UTR with regulatory sequences from the *cdc2* or *adh1* genes does not affect size dependence of Cdc13 expression (Figure 3A). Together, these results demonstrate that size dependence is encoded in the *cdc13* ORF.

**Figure 3.**
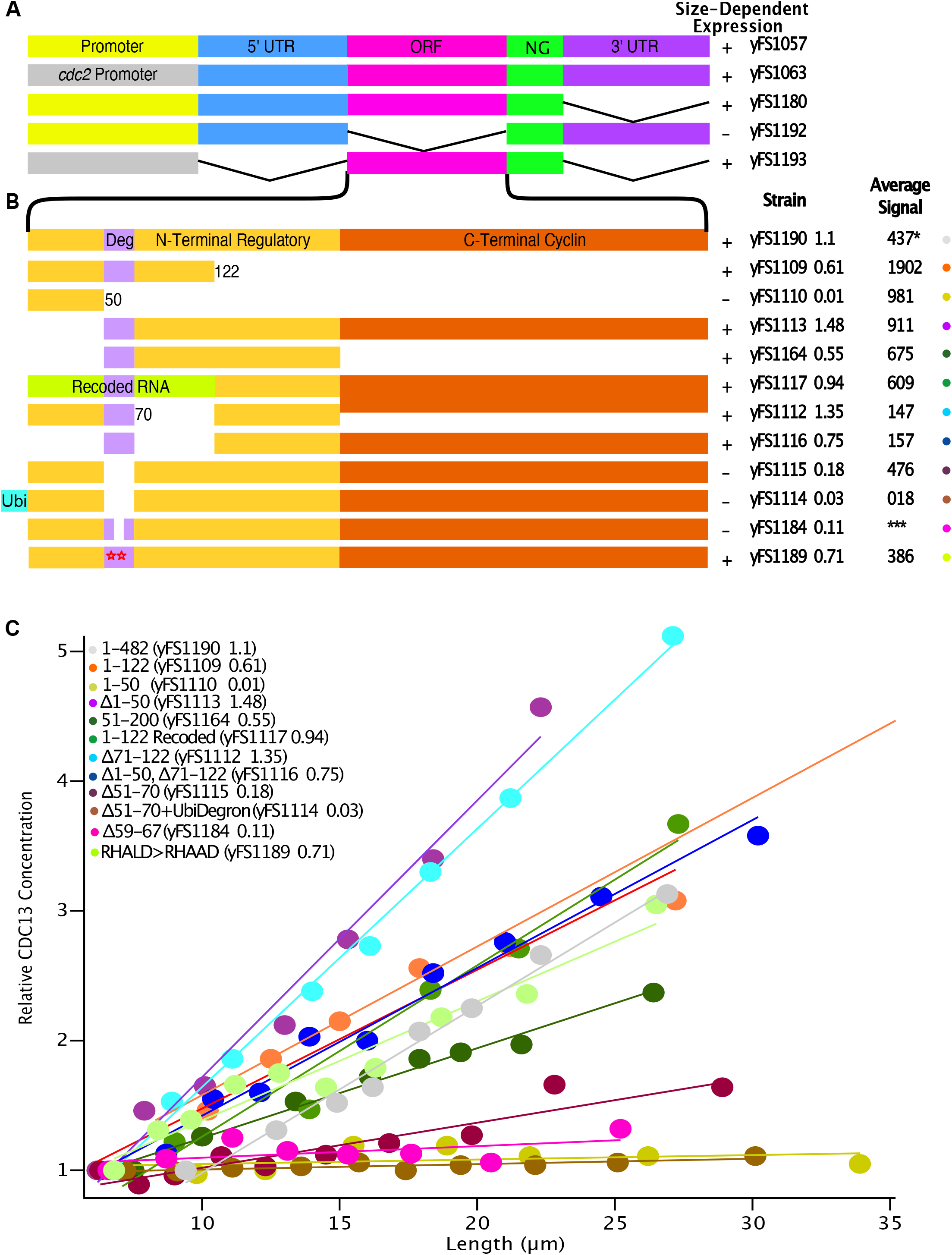
Cdc13 Size-Dependent Regulation is Encoded in a 20 Amino Acid N-terminal Motif. A) Cdc13 Size-Dependent Regulation is Encoded in Its ORF. The size dependence of Cdc13 expressed from various depicted constructs was assessed as in Figure 1A. B) Deletion Analysis of the Cdc13 ORF. The size dependence of Cdc13 expressed from various ORF deletions was assessed as in Figure 1A, cell size was manipulated by growing *ZEV:wee1* cells in 0 to 100 nM beta-estradiol. C) Data from the Deletion Analysis of the Cdc13 ORF. The data from B is plotted as the average values of 50-cell bins. The legend states the strain name and the correlation metric for the data. *This sample was imaged on both the DeltaVision and MICA microscopes. Its average intensity was calculated only for the dataset acquired using the DeltaVision, which was used for all other constructs. *** No averages are given because this sample was imaged on the MICA.

To further localize the sequence required for size-dependent expression within the *cdc13* ORF, we performed a deletion analysis. To be able to quantitatively distinguish size-dependent from size-independent expression, we developed a metric that represents the correlation between size and relative protein concentration. The metric, which is effectively the slope of a linear fit to the data on a relative size to relative concentration plot, is 1 if Cdc13 concentration is perfectly correlated with size and is 0 if it is uncorrelated with size. In practice, we find a clear distinction between size-dependent expression (with a score of greater than 0.5) and size-independent expression (with a score less than 0.5) (Figure 3C).

We initially split the *cdc13* ORF in the two parts that encode its N-terminal unstructured regulatory domain and its C-terminal cyclin domain (Chethan et al., 2025), and found that the N-terminal region is necessary for size-dependent expression (Figure 3B). Further deletions narrowed the critical region down to a 20-codon region that includes the D-box degron, which is responsible for the APC-dependent degradation of Cdc13 during late mitosis and G1.

Finding that a small region encoding the N-terminus of Cdc13 is required for its size-dependent expression could indicate that size-dependence is conferred by the amino acids in that part of the protein or by the RNA that encodes them. To distinguish between these two possibilities, we recoded the first 122 codons of *cdc13*, replacing every ambiguous nucleotide (123/366) to change the RNA sequence as much as possible without changing the encoded protein sequence. Such recoding does not affect the size-dependent expression of Cdc13 (Figure 3C), suggesting that size-dependence is encoded in Cdc13’s amino acids, not the nucleotides of its transcript.

Constructs that retain the degron region retain size-dependent expression. However, it is more difficult to determine if constructs that lack this region are expressed in a size-dependent manner because they are not degraded during mitosis. Therefore, the amount of Cdc13 in such cells may reflect Cdc13 inherited from the parental cell, in addition to newly synthesized Cdc13, obscuring the kinetics of Cdc13 synthesis. This complication is manifest in the fact that a construct expressing only the N-terminal 50 amino acids of Cdc13 shows no size dependence (correlation metric = 0.01) and has an overall signal that is more than two-fold higher than the full-length construct (Figure 3B). This overexpression of Cdc13 during mitosis would plausible be lethal, due to interference with mitotic exit (Murray et al., 1989; Yamano et al., 1996), and may explain why we have been unable to create strains carrying constructs that would express the C-terminal cyclin domain without the N-terminal regulatory domain.

Nonetheless, we were able to create a strain that expresses a Cdc13 allele lacking just amino acids 51-70 (Cdc13Δ51-70), which contain the D-box degron. This construct is expressed at levels comparable to the full-length level and is not toxic. From these results, we surmise that the rest of the N-terminal regulatory region confers instability to Cdc13, relative to that of its C-terminal cyclin domain. Cdc13Δ51-70 is expressed in a size-independent manner, demonstrating that those 20 amino acids are necessary for size-dependent expression. Additionally, it is not as efficiently degraded during mitosis, as binucleate cells show a three-fold increase in average mean fluorescence compared to binucleate cells expressing full-length Cdc13 (Figure S3). To confirm that the increased mitotic stability of Cdc13Δ51-70 is not obscuring size-dependent expression, we fused a ubiquitin degron to the N-terminus of the protein. The N-terminal ubiquitin is expected to be removed by constitutive ubiquitin hydrolases, exposing an N-terminal tyrosine that destabilizes the protein (Houser et al., 2012). This ubiquitin degron decreases the average fluorescence intensity of cells carrying Cdc13Δ51-70 about tenfold, but does not affect its size-independence. In addition, we created a construct in which the N-terminal region flanking amino acids 51-70 was deleted and found that this construct was expressed in a size-dependent manner (Figure 3B, C). Therefore, we conclude that the 20 amino acids that include the D-box degron are necessary and sufficient within the Cdc13 N-terminus for size dependent expression.

To further investigate the role of the 20 amino acids from positions 51 to 70, we introduced additional deletions in this region. We found that deletion of amino acids 59-67, which corresponds to the sequence for the minimal destruction box (RHALDDVSN)(Yamano et al., 1996), renders Cdc13 expression size-independent (Figure 3C). To more specifically test the degron function of this sequence, we mutated the RHALD degron-consensus sequence to AHAAD to disable APC-dependent degradation (Yamano et al., 1996) and found that Cdc13 continues to be expressed in a size-dependent manner (Figure 3C). We then directly tested the role of the APC in the size-dependence of Cdc13 expression. We arrested cell in G2, using the *cdc2-M17as* analog-sensitive allele and inactivated the APC, using the *cut4-533ts* temperature-sensitive allele. In this situation, Cdc13 expression is still size dependent (Figure S4). Therefore, we conclude that APC-dependent degradation is not required for size-dependent expression of Cdc13.

### Size-Dependent Expression of Cdc13 is not Necessary for G2 Size Control in Fission Yeast

Based on the correlation between concentration and cell size for Cdc13 (Creanor and Mitchison, 1996; Patterson et al., 2019; Curran et al., 2022) it has been proposed to be involved in fission yeast G2 size control. Our structure-function analysis of Cdc13 identified deletions that are expressed at size-independent concentrations (Figure 3). In particular, it seemed plausible that Cdc13-Δ51-70, which is expressed in a size-independent manner and lacks only 20 amino acids from its N-terminal regulatory domain, would be a functional allele. To test this possibility, we deleted the endogenous *cdc13* in strains carrying both *cdc13-Δ51-70* and *ubi-cdc13-Δ51-70*. In both cases, we obtained strains that are healthy, grow at a normal rate and maintain robust size homeostasis (Figure 4). The single mutant strains are longer than the wild type (Figure 4), indicating that the size-independent alleles of *cdc13* have reduced activity. To assay this allele at an endogenous context, we deleted the codons encoding these 20 amino acids from the endogenous *cdc13* locus using Cas9-directed gene conversion (Torres-Garcia et al., 2020). We obtained strains that are healthy and maintain robust size homeostasis, with an average length at septation closer to the wild type compared with the other two mutants (Figure 4A). One plausible explanation is that Cdc13 expression in this strain is driven from its own promoter, whereas in the other two mutants, expression is driven by the *cdc2* promoter. Importantly, the majority of mutant cells actually divide in a narrow range, confirming robust size homeostasis (Figure 4B). In any case, size-dependent expression of Cdc13 does not appear to be required for G2 size control.

**Figure 4.**
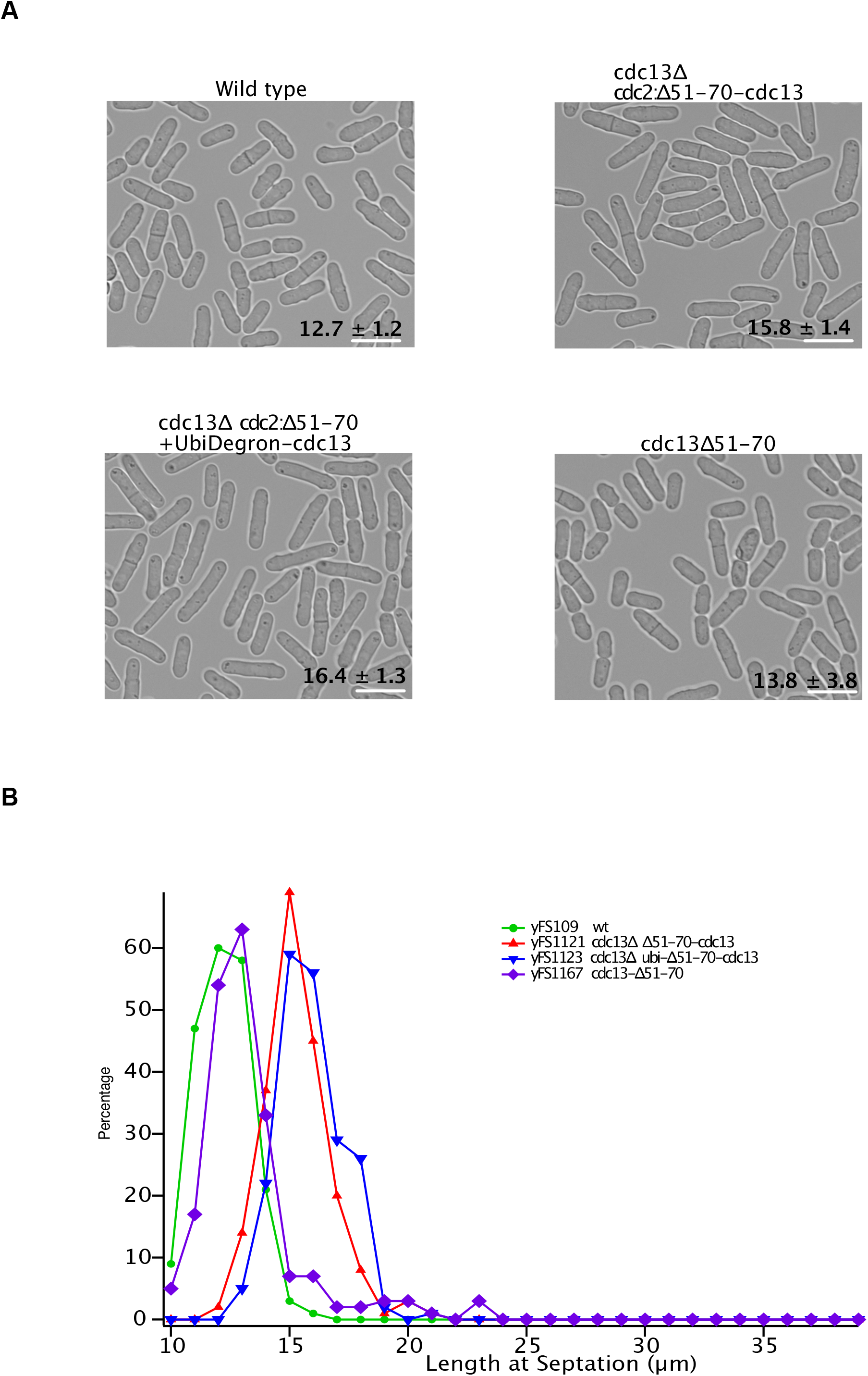
Cdc13 is not Required for Size Control. A) Size-Dependent Expression of Cdc13 is not Required for Viability. Wild-type (yFS110), *cdc13Δ cdc13-ubi-Δ51-70* (yFS1123), *cdc13-Δ cdc13-Δ51-70 (yFS1121), and cdc13-Δ51-70* (yFS1167) cells were grown to mid-log and photographed. Magnification 40x; Scale bar, 10μm. B) Size-Dependent Expression of Cdc13 is not Required for Size Homeostasis. The distribution of lengths at septation for the cultures shown in A. n = 200 for each strain.

## Discussion

We have characterized the size dependence of the fission yeast B-type cycle, Cdc13, and investigated its role in size-dependent regulation of the G2/M transition. We find that Cdc13 is expressed in a size-dependent manner and that it’s size dependence is conferred post-transcriptionally. However, we find that size-dependent expression of Cdc13 is not required for size control of the G2/M transition.

We conclude that Cdc13 is expressed in a size-dependent manner, and specifically that the correlation between its concentration and cell size is not an indirect result of time-dependent expression. By varying the size at division in asynchronous cultures through the manipulation of Wee1 kinase activity, we obtained cultures that begin the cell cycle at various sizes. If Cdc13 is expressed in a time-dependent manner, it should be expressed at the same level in all cells at the beginning of the cell cycle, independent of cell size. If, on the other hand, it is expressed in a size-dependent manner, it should exhibit similar concentrations at similar sizes, independent of how long the cells have been in the cell cycle. We observe the latter result, with cells that are 15 μm long have about the same concentration of Cdc13 regardless of whether they are at the beginning, the middle, or the end of their cell cycle (Figure 1B). A recent report describes a similar experiment using deletion of *cdr2*, which encodes a negative regulator of Wee1, to affect cell size, but reaches the opposite conclusion (Miller et al., 2022). We have no clear explanation of the discrepancy between these two sets of experiments. However, we will note the size differential at division in the experiments reported in the Miller paper is 34%, whereas the size differential of the experiments presented in Figure 1B is 162%, making the distinction between size- and time-dependent behavior more robust in our experiments. Nonetheless, Miller et al. present other data in support of their conclusion that Cdc13 is expressed in a time-dependent manner, for which we have no explanation.

To investigate how Cdc13 is regulated in a size-dependent manner, we narrowed down the sequence necessary and sufficient for size-dependent expression to a 20-codon motif in the region encoding its N-terminal regulatory domain (Figure 3). To determine if the RNA sequence of this motif, or the protein sequence that it encodes, is responsible for size-dependent expression, we recoded the first 122 codons of Cdc13, replacing every ambiguous nucleotide with one that would not change the encoded protein sequence, which changed one third of the RNA sequence. With this recoding, we hoped to disrupt any regulatory motifs or secondary structures contained in the RNA sequence without affecting the encoded protein. The fact that the recoded sequence is expressed in a size-dependent manner leads us to conclude that size dependence is conferred by the protein sequence, although it is possible that the recoding failed to disrupt a critical RNA regulatory sequence.

The 20-amino-acid motif that we identified as being necessary and sufficient for size-dependence contains the D-box motif recognized by the APC for degradation of Cdc13 during post-anaphase mitosis and G1. It is not expected that the APC would regulate Cdc13 stability during G2 because the APC is not thought to be active then. Consistent with this expectation, point mutations in the D-box sequence that inactivate its mitotic-degron function do not affect the size-dependence of Cdc13. We further tested this expectation by expressing Cdc13-NG in a *cut4-533ts* allele background and found that inactivation of this essential APC subunit does not interfere with the size-dependent expression of Cdc13 (Figure S4). Therefore, we conclude that APC is not involved in Cdc13’s size-dependent expression.

It is surprising that Cdc13Δ51-70, which lacks its primary APC degron, is viable. Removal of the APC degron from cyclin B in budding yeast and *Xenopus* extracts causes anaphase arrest with high CDK levels (Murray et al., 1989; Surana et al., 1993; Sigrist et al., 1995). The fact that we are unable to create strains carrying just the C-terminal cyclin domain of Cdc13 suggests that overexpression of a stable Cdc13 is also lethal. The viability of strains expressing Cdc13Δ51-70 may be due to a secondary APC degron that has been identified in the very N-terminus of Cdc13 (Yamano et al., 1996) or to the ability of Wee1 to inhibit CDK post anaphase (Lianga et al., 2013); either or both of these mechanisms may allow sufficient reduction in CDK activity to allow cells to enter G1.

The fact that a size-dependent amount of Cdc13 is expressed from a size-independent amount of *cdc13* mRNA suggests that Cdc13 is regulated by size dependent translation or degradation. Since Cdc13 appears to be regulated by an internal protein motif, it is unlikely to be regulated by size-dependent translational initiation. Even if its regulation affects elongation, the involvement of an internal protein motif is unexpected. Therefore, we speculate that Cdc13 abundance is regulated by size-dependent protein turnover, but investigation of that hypothesis will require further investigation.

Regardless of the mechanism of Cdc13’s expression, how any such mechanism could be size-dependent is hard to conceive. To the extent that proteins and RNA generally maintain a constant concentration as cells grow (Zhurinsky et al., 2010; Marguerat et al., 2012; Schmoller and Skotheim, 2015), there should be no difference in translation (or any other biochemistry) between small and large cells. The one thing that does universally change with cell size is the concentration of DNA, with the ratio of protein to DNA increasing as cells grow. This change has been proposed as the basis for a number of different mechanisms by which transcription could be made to be size dependent (Wang et al., 2009; Schmoller and Skotheim, 2015; Schmoller et al., 2015; Dorsey et al., 2018). Therefore, we speculate that it is unlikely that Cdc13’s synthesis or degradation is directly size-dependent. Instead, we offer the hypothesis that Cdc13’s translation is regulated by a size-dependent regulator that is itself regulated by size-dependent transcription.

Cdc13’s size-dependent expression makes it a prime candidate for a central cell-size regulator as an accumulating activator of the G2/M transition in fission yeast (Figure 1A, C and Patterson et al., 2019; Creanor and Mitchison, 1996; Curran et al., 2022). Our analysis, which identified Cdc13Δ51-70 as a functional allele that is expressed in a size-independent manner, allowed us to test that hypothesis directly. The fact that cells expressing Cdc13Δ51-70 as their only copy of Cdc13 are viable, healthy and maintain size homeostasis demonstrates that size-dependent expression of Cdc13 is not required for cell-size control (Figure 4A).

The fact that Cdc13 is not required for G2/M size control in fission yeast does not mean that it is not involved in the process. Cdc25, the other mitotic activator that accumulates in a size-dependent manner (Figure 4A and Keifenheim et al., 2017; Creanor and Mitchison, 1996; Moreno et al., 1990; Curran et al., 2022; Miller et al., 2022), is likewise a candidate for a central cell-size regulator as an accumulating activator of the G2/M transition in fission yeast. Similarly, Cdr2 has been proposed to regulate G2/M size control in fission yeast, via its regulation of Wee1 (Pan et al., 2014; Facchetti et al., 2019; Miller et al., 2022), although it too has been shown to be dispensable for such regulation (Facchetti et al., 2019; Miller et al., 2022). It has been proposed that all three—Cdc13, Cdc25 and Cdr2—could be redundantly required for G2/M size control in fission yeast (Miller et al., 2022). Alternatively, it is possible that many positive and negative regulators of the G2/M transition collaborate to regulate size control (Chen et al., 2020). Either way, a detailed understanding of the mechanisms underlying size-dependent expression of cell-cycle regulators will be crucial for elucidating the strategies that cells use to maintain stable cell size.

## Methods

### Strain and plasmid construction

Strains and plasmids were created using standard approaches (Forsburg and Rhind, 2006). Cultures were grown in YES at 30°C, unless otherwise noted. Strains, plasmids and primers used in this study are listed in Tables 1, 2 and 3. Plasmids were created using Gibson Assembly (NEB) as detailed in Table 4.

**Table 1:**
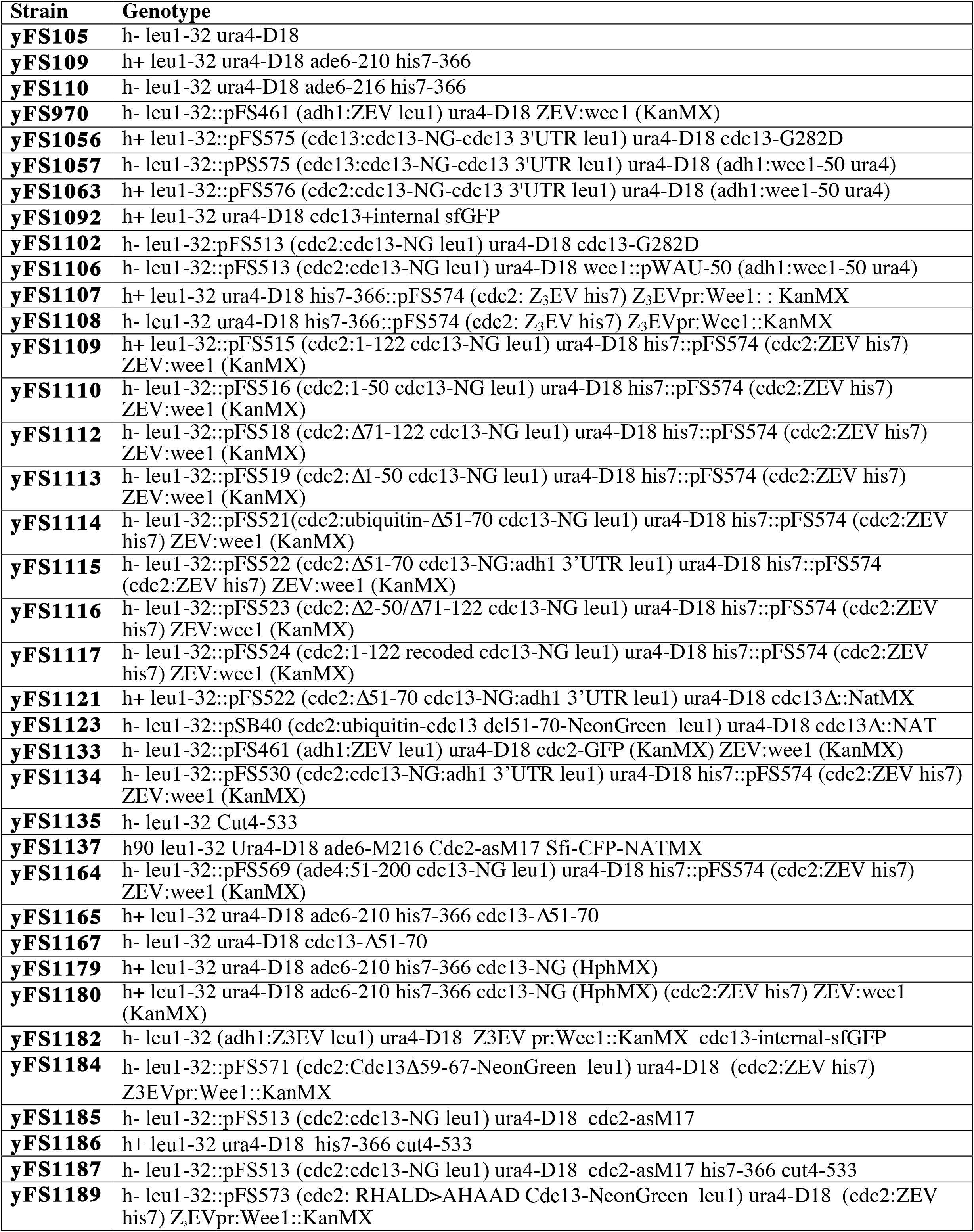
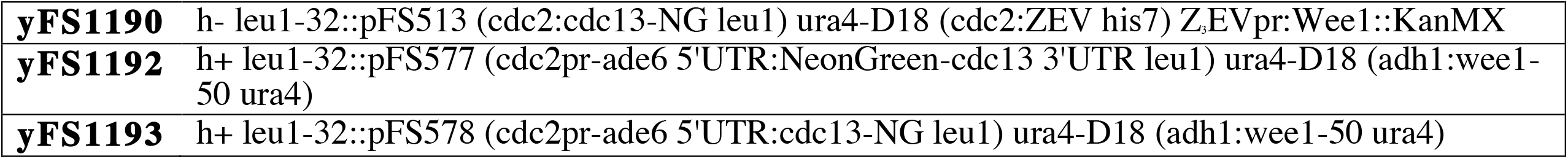
Strains used.

**Table 2:**
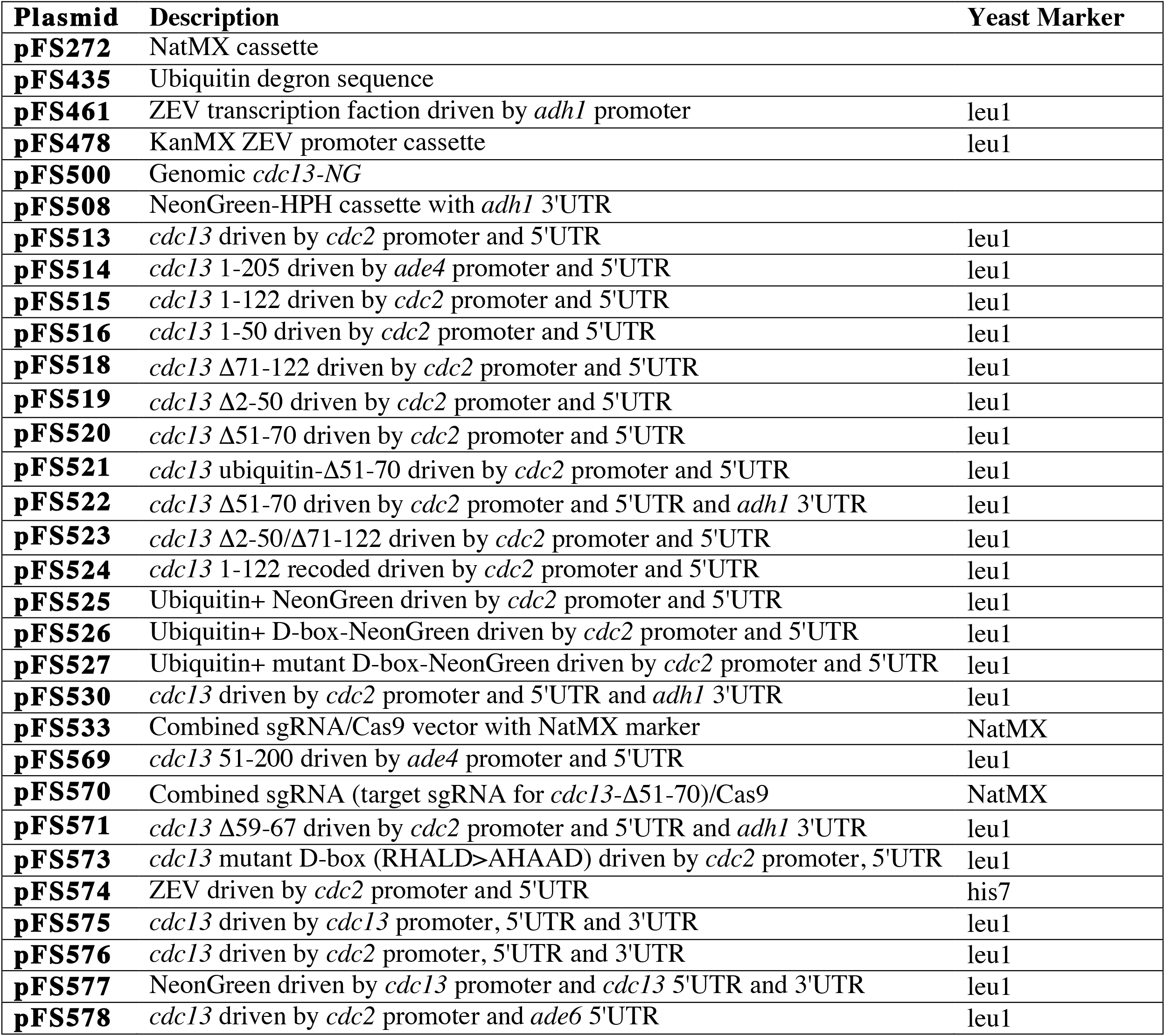
Plasmids used.

| **Plasmid** | **Description** | **Yeast Marker** |
| --- | --- | --- |
| **pFS272** | NatMX cassette |  |
| **pFS435** | Ubiquitin degron sequence |  |
| **pFS461** | ZEV transcription faction driven by *adh1* promoter | leu1 |
| **pFS478** | KanMX ZEV promoter cassette | leu1 |
| **pFS500** | Genomic *cdc13-NG* |  |
| **pFS508** | NeonGreen-HPH cassette with *adh1* 3'UTR |  |
| **pFS513** | *cdc13* driven by *cdc2* promoter and 5'UTR | leu1 |
| **pFS514** | *cdc13* 1-205 driven by *ade4* promoter and 5'UTR | leu1 |
| **pFS515** | *cdc13* 1-122 driven by *cdc2* promoter and 5'UTR | leu1 |
| **pFS516** | *cdc13* 1-50 driven by *cdc2* promoter and 5'UTR | leu1 |
| **pFS518** | *cdc13* Δ71-122 driven by *cdc2* promoter and 5'UTR | leu1 |
| **pFS519** | *cdc13* Δ2-50 driven by *cdc2* promoter and 5'UTR | leu1 |
| **pFS520** | *cdc13* Δ51-70 driven by *cdc2* promoter and 5'UTR | leu1 |
| **pFS521** | *cdc13* ubiquitin-Δ51-70 driven by *cdc2* promoter and 5'UTR | leu1 |
| **pFS522** | *cdc13* Δ51-70 driven by *cdc2* promoter and 5'UTR and *adh1* 3'UTR | leu1 |
| **pFS523** | *cdc13* Δ2-50/Δ71-122 driven by *cdc2* promoter and 5'UTR | leu1 |
| **pFS524** | *cdc13* 1-122 recoded driven by *cdc2* promoter and 5'UTR | leu1 |
| **pFS525** | Ubiquitin+ NeonGreen driven by *cdc2* promoter and 5'UTR | leu1 |
| **pFS526** | Ubiquitin+ D-box-NeonGreen driven by *cdc2* promoter and 5'UTR | leu1 |
| **pFS527** | Ubiquitin+ mutant D-box-NeonGreen driven by *cdc2* promoter and 5'UTR | leu1 |
| **pFS530** | *cdc13* driven by *cdc2* promoter and 5'UTR and *adh1* 3'UTR | leu1 |
| **pFS533** | Combined sgRNA/Cas9 vector with NatMX marker | NatMX |
| **pFS569** | *cdc13* 51-200 driven by *ade4* promoter and 5'UTR | leu1 |
| **pFS570** | Combined sgRNA (target sgRNA for *cdc13*-Δ51-70)/Cas9 | NatMX |
| **pFS571** | *cdc13* Δ59-67 driven by *cdc2* promoter and 5'UTR and *adh1* 3'UTR | leu1 |
| **pFS573** | *cdc13* mutant D-box (RHALD>AHAAD) driven by *cdc2* promoter, 5'UTR | leu1 |
| **pFS574** | ZEV driven by *cdc2* promoter and 5'UTR | his7 |
| **pFS575** | *cdc13* driven by *cdc13* promoter, 5'UTR and 3'UTR | leu1 |
| **pFS576** | *cdc13* driven by *cdc2* promoter, 5'UTR and 3'UTR | leu1 |
| **pFS577** | NeonGreen driven by *cdc13* promoter and *cdc13* 5'UTR and 3'UTR | leu1 |
| **pFS578** | *cdc13* driven by *cdc2* promoter and *ade6* 5'UTR | leu1 |

**Table 3:**
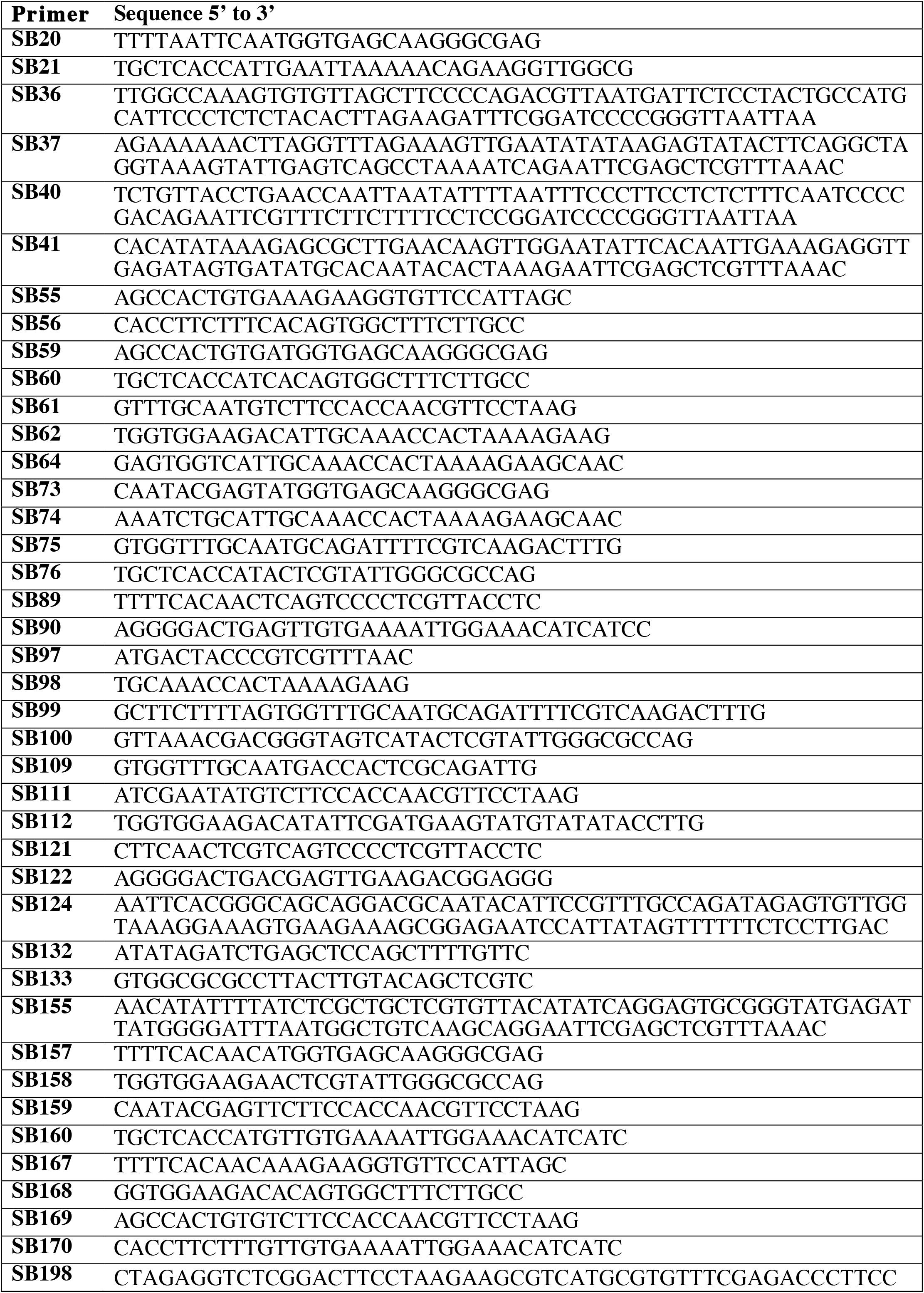
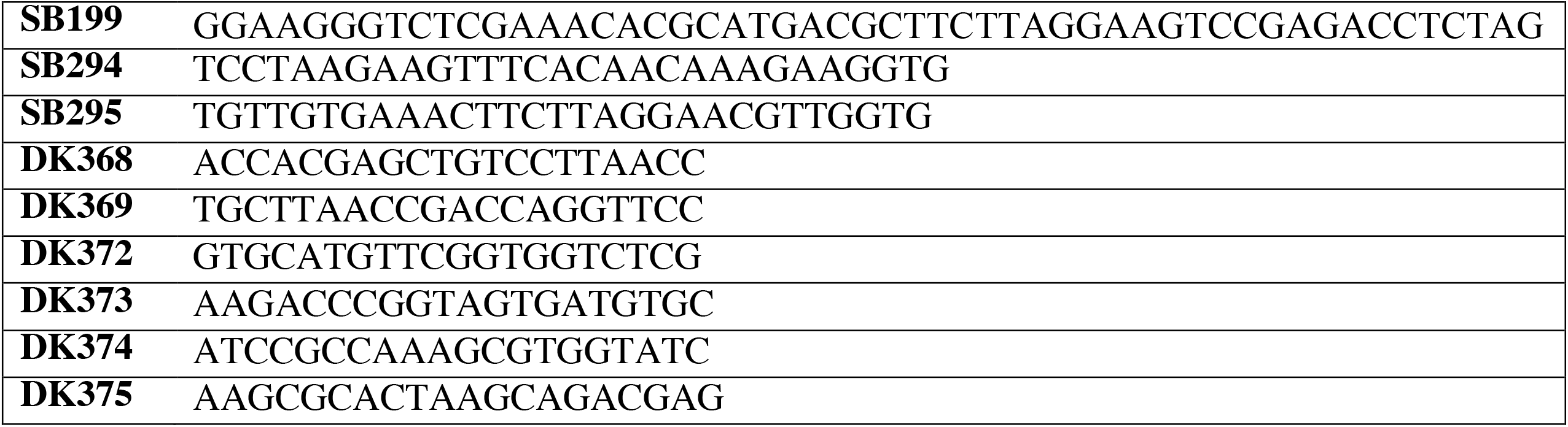
Primers Used.

**Table 4:**
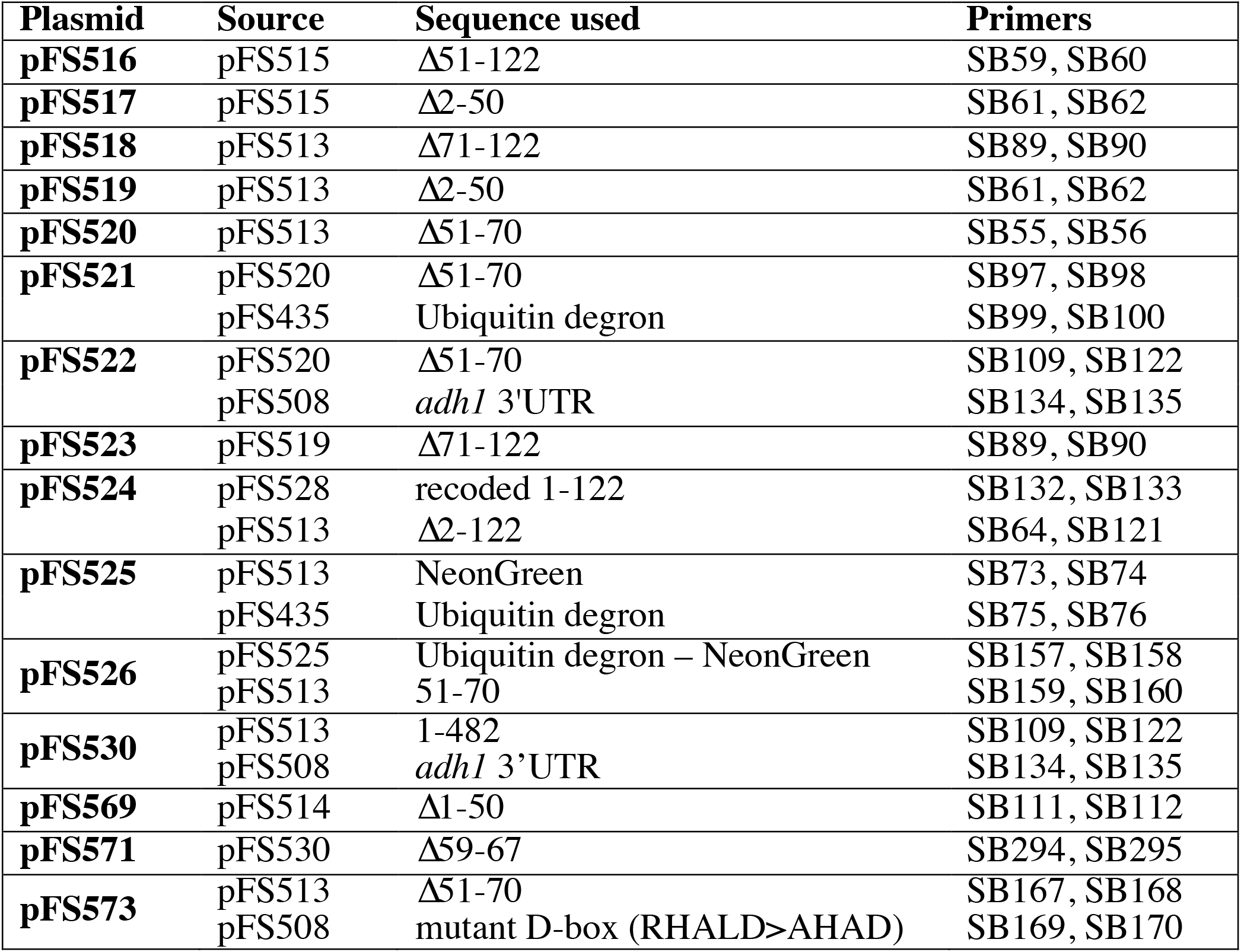
Plasmid Construction Details.

To create a recoded version for *cdc13* in pFS524, the following sequence, designed to replace every ambiguous nucleotide without changing the encoded amino acids, in which the replaced nucleotides are lowercase and codons 51-70 are italic, was purchased from Genewiz (Azenta Life Sciences).

ATGACcACtCGcaGaTTgACcCGtCAaCAtCTgTTaGCgAAcACtTTaGGtAAtAAcGAtGAgAAcCAc CCcTCtAAtCAcATcGCtaGaGCgAAgAGtTCcTTaCAtTCcTCgGAgAAcTCcTTgGTgAAcGGtAAa AAAGCtACcGTa*TCcTCtACtAAtGTcCCcAAaAAaCGcCAcGCaTTaGAcGAcGTcTCtAAcTTcCAt AAt*AAgGAgGGcGTgCCcTTgGCgAGcAAgAAtACgAAcGTtAGgCAtACcACaGCgTCcGTgAGcACt CGcaGaGCgCTgGAaGAgAAaTCaATcATtCCcGCgACtGAcGAcGAgCCaGCcTCtAAaAAaCGcCGtCAgCCcTCcGTcTTcAAcTCg

To generate the *cdc13-Δ51-70* target sgRNA/Cas9 construct (pFS570), primers SB198 and SB190 were annealed and cloned into pFS533 using the NEB Golden Gate Assembly Kit (BsaI-HF v2) following the protocol as described (Torres-Garcia et al., 2020).

The *cdc13Δ-51-70* strain (yFS1167) was constructed by using the Cas9 method as previously described (Torres-Garcia et al., 2020). A single guide RNA targeting amino acids 51-70 of Cdc13 (5’-TCCTAAGAAGCGTCATGCGT-3’) was used. The homologous recombination (HR) repair template was generated by PCR amplification from the plasmid pFS522 using primers SB200 and SB201; the sequence of the HR template follows.

TAACTCGCCAGCACCTATTGGCAAATACCTTGGGCAACAATGACGAAAATCATCCTTCAAACCATATTGCCCGTGCAAAAAGCTCTTTGCACTCTTCAGAAAATTCTTTAGTAAATGGCAAGAAAGCCACTGTGAAAGAAGGTGTTCCATTAGCTAGTAAAAACACAAATGTCAGACACACTACTGCTTCTGTCAGTACCCGTCGTGCTCTCGAGGAAAAGTCTATAATCCCTGCAACAGATGATGAACCCGCTTCCAAGAAGCGTCGCCAACCTTCTGTTTTT

### Fluorescent Microscopy

Cells were grown in YES to mid log phase. To obtain different sized cells, cells were either grown at different temperatures or in presence of different levels of beta-estradiol, as appropriate. Cells were fixed in 100% methanol at -80°C for at least 15 minutes or up to 3 weeks. Cells were rehydrated in 1X PBS and imaged by widefield fluorescence microscopy with a DeltaVision-enabled Olympus IX71 inverted microscope with a 60x/1.42 oil-immersion objective or a Leica MICA. Images acquired were analyzed using ImageJ (Schneider et al., 2012) and pomBseen (Ohira and Rhind, 2022). Binucleate cells, which comprise the post-anaphase cells in the population, were excluded from analysis because Cdc13 is degraded from anaphase through G1. For the quantification of Cdc13 size dependence, we divided the relative concentration by the relative size. Values were baseline-corrected by subtracting 1 from both measurements before calculating the ratio.

### Single Molecule RNA Fluorescence In Situ Hybridization (smFISH)

smFISH samples were prepared according to a modification of published protocols (Trcek et al., 2012; Schulz et al., 2013). Briefly, cells were fixed in 4% formaldehyde and the cell wall was partially digested using Zymolyase. Cells were permeabilized in 70% EtOH, pre-blocked in BSA and salmon sperm DNA, and incubated over-night with custom Stellaris oligonucleotides sets (Biosearch Technologies) designed against cdc13 (CAL Fluor Red 610) and rpb1 (Quasar 670) mRNAs (Table S2). Cells were mounted in ProLong Gold antifade reagent with DAPI (Life Technologies) and imaged on a Leica TCS Sp8 confocal microscope, using a 63x/1.40 oil-immersion objective. Optical z sections were acquired (z-step size 0.3 microns) for each scan to cover the depth of the cells. Cell boundaries were outlined manually and single mRNA molecules were identified and counted using the FISH-quant MATLAB package (Mueller et al., 2013). Cell area, length and width were quantified using custom ImageJ macros. The FISH-quant detection technical error was estimated at 6%–7% by quantifying rpb1 mRNAs simultaneously with two sets of probes labeled with different dyes.

## Acknowledgements

We are grateful to Christiana Baer and the Sanderson Center for Optical Experimentation (SCOpE) for access to, and technical support for, the DeltaVision and MICA microscopes. Strains for this work were obtained from the National BioResource Project/Yeast Genetic Resource Center (NBRP/YGRC). We are grateful to Silke Hauf for sharing strains. This work was supported by NIGMS R01 GM134300 and R35 GM148359 to NR.

---

**Figure S1.**
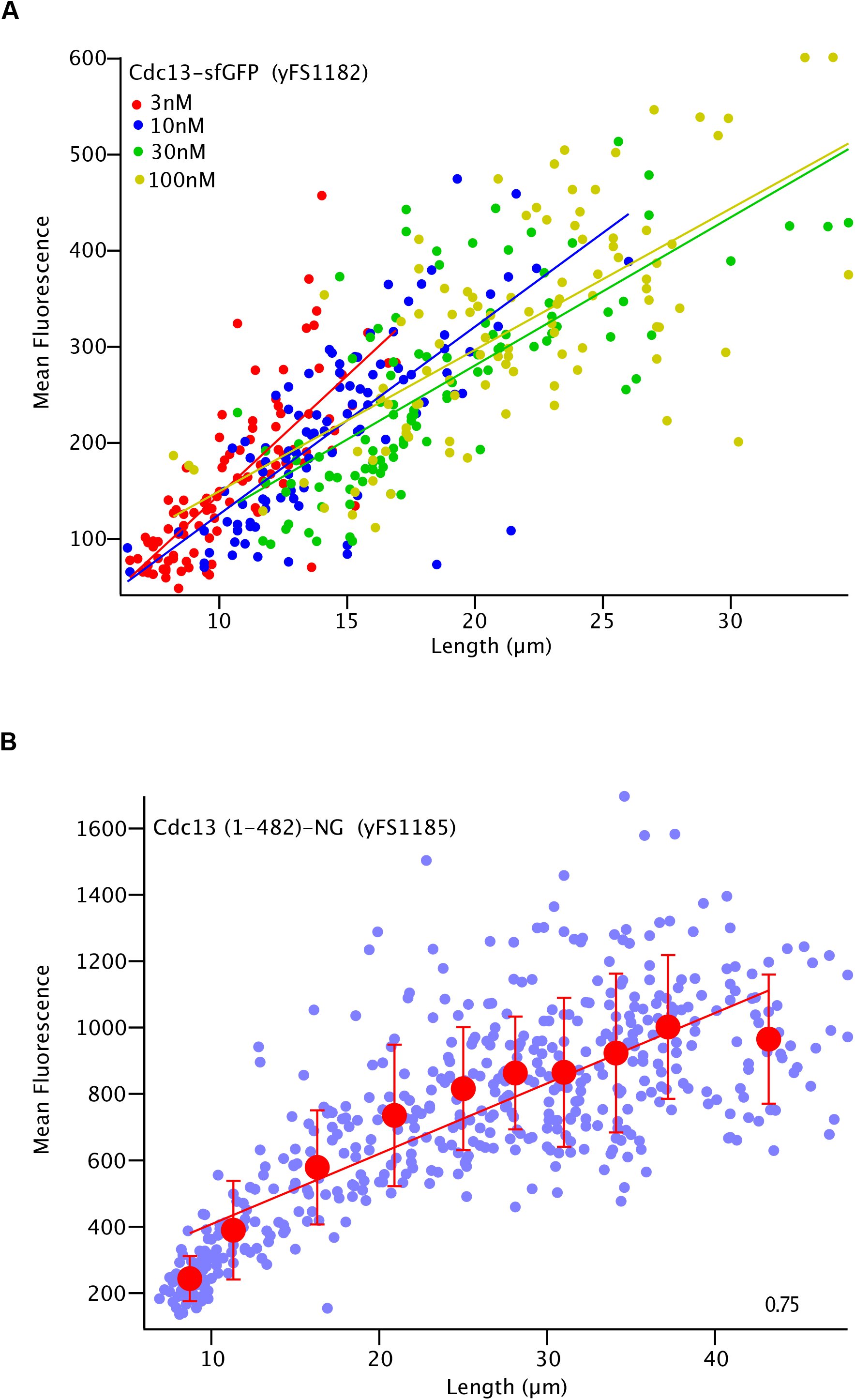
Cdc13 is Expressed in a Size-Dependent Manner. A) Asynchronous cells expressing Cdc13-sfGFP from its endogenous locus (yFS1182) were analyzed by widefield fluorescence microscopy. Asynchronous cultures of cells spanning various size ranges were obtained by growth in 3nM, 10nM, 30nM, and 100nM estradiol to regulate the activity of Wee1 expressed from the *ZEV* promoter. Data from different cultures is plotted in different colors. B) *cdc2asM17 cdc2:cdc13-NeonGreen (*yFS1185) cells were treated with 5μM 1NM-PP1 to arrest cells at G2 and collected at time points at 0, 2, 4, 5, and 6 hours and analyzed by widefield fluorescence microscopy.

**Figure S2.**
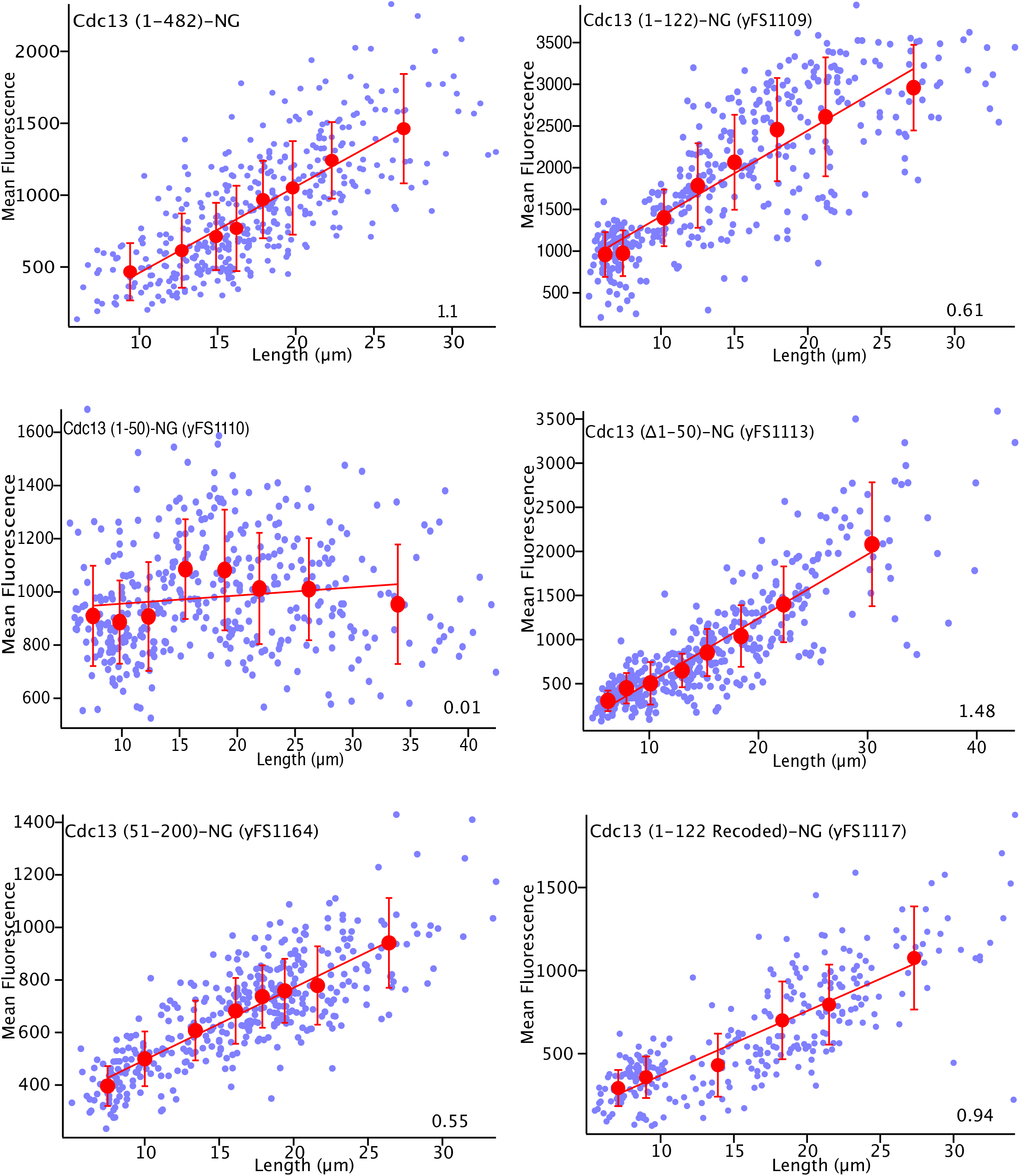
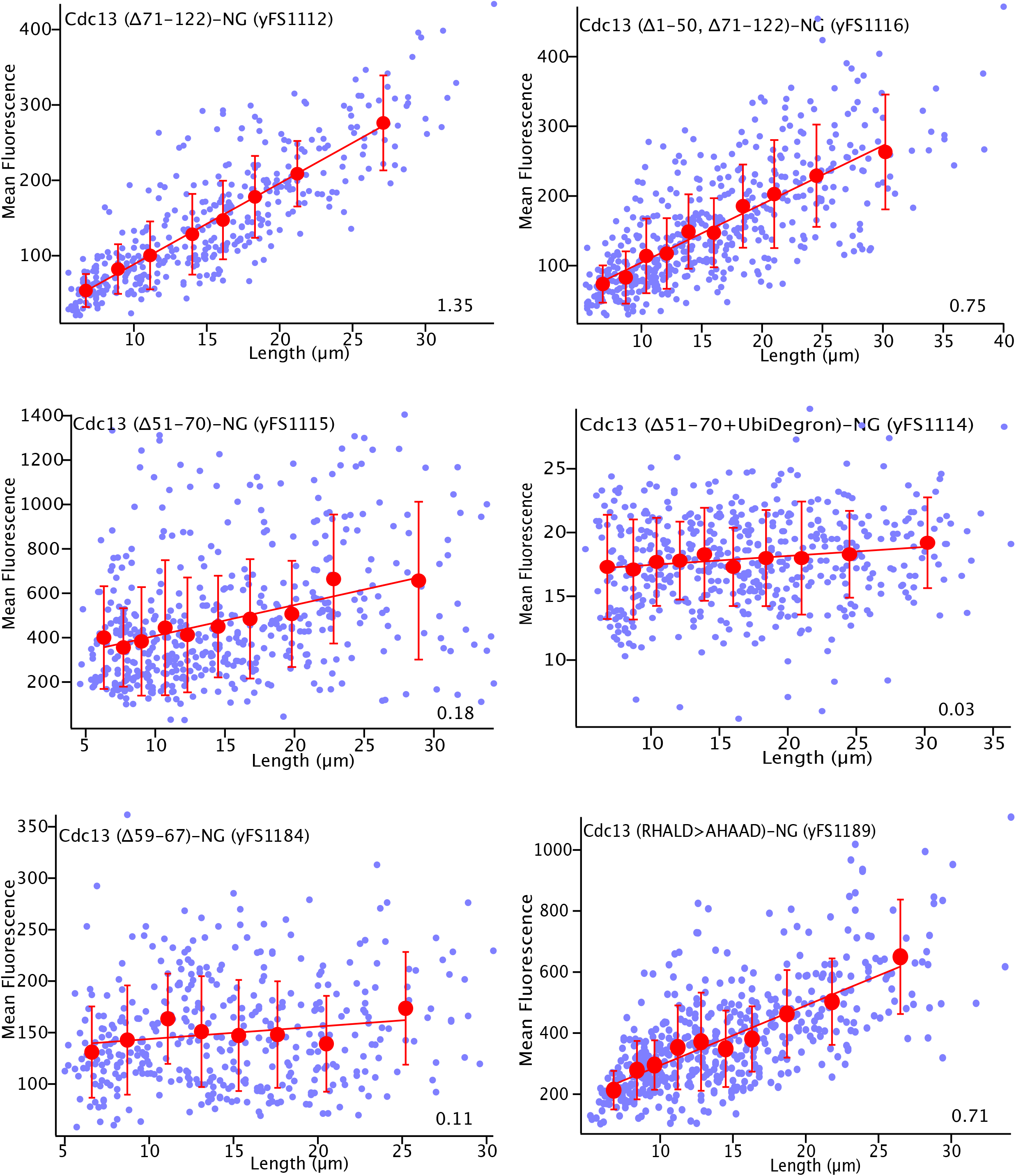
Cdc13 Size-Dependent Regulation is Encoded in a 20 Amino Acid N-terminal Motif. The size dependence of Cdc13 expressed from various ORF deletions was assessed as in Figure 1A; cell size was manipulated by growing *ZEV:wee1* cells in 0 to 100 mM beta-estradiol. The small symbols represent individual cells; the large symbols represent an average of the data in 50-cell bins. These data correspond to each *cdc13* ORF deletion construct presented in Figure 3C.

**Figure S3.**
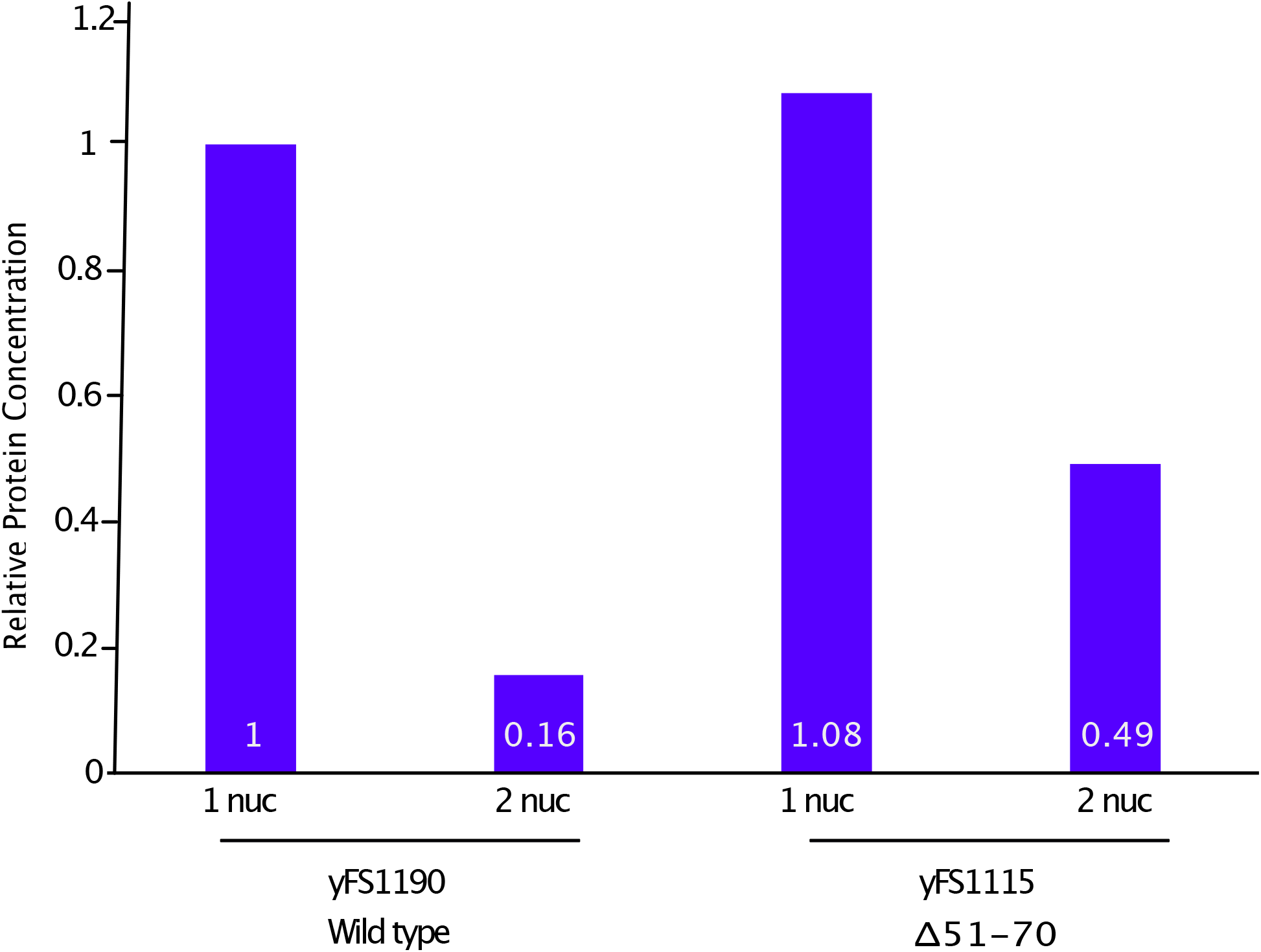
Cdc13 Lacking D-box is not Properly Degraded During Mitosis. Comparison of the average mean fluorescence between mono-nucleate and bi-nucleate cells in strains *cdc2:cdc13-NeonGreen (*yFS1190) and *cdc2:cdc13-Δ51-70*-NeonGreen (yFS1115).

**Figure S4.**
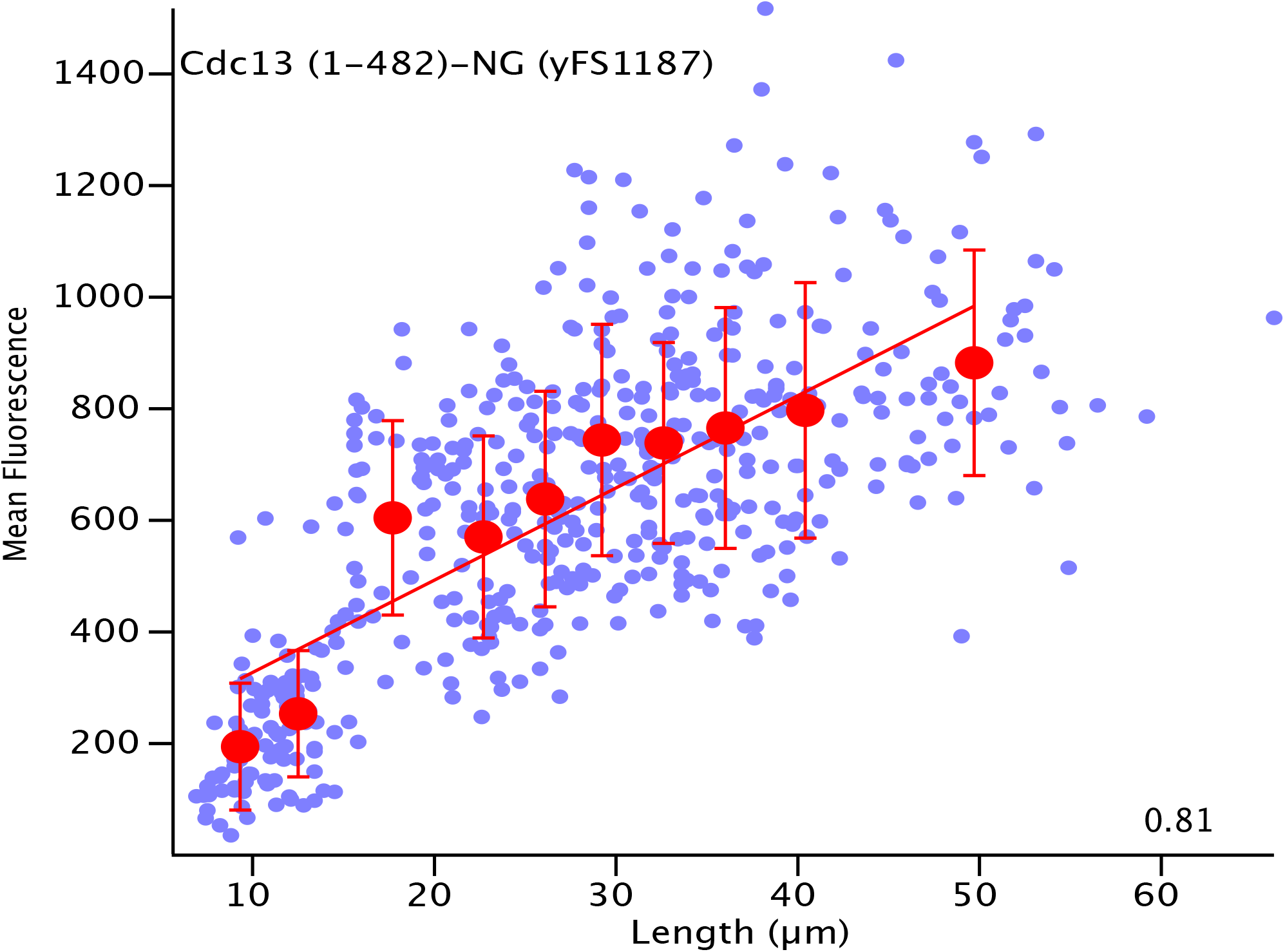
A Mutation in an Essential APC Subunit does not Interfere with the Size-Dependent Expression of Cdc13. *cdc2asM17 cdc2:cdc13-NeonGreen cut4-533* (yFS1187) was analyzed as in Figure S1B, except that cells were shifted to the non-permissive temperature of 35 °C at time 0.

## References

Amodeo, A. A., and Skotheim, J. M. (2016). Cell-Size Control. Cold Spring Harb Perspect Biol 8, a019083.

Aoi, Y., Kawashima, S. A., Simanis, V., Yamamoto, M., and Sato, M. (2014). Optimization of the analogue-sensitive Cdc2/Cdk1 mutant by in vivo selection eliminates physiological limitations to its use in cell cycle analysis. Open Biol 4, 140063.

Bandyopadhyay, S., Ghosh, P. M., Basu, S. et al. (2017). Antagonistic regulation of cyclin expression by the bZIP transcription factors Pcr1 and Atf1 during G2/M transition. FEMS Microbiology Letters 364,

Blanco, M. A., Sánchez-Díaz, A., de Prada, J. M., and Moreno, S. (2000). APC(ste9/srw1) promotes degradation of mitotic cyclins in G(1) and is inhibited by cdc2 phosphorylation. EMBO J 19, 3945–3955.

Booher, R. N., Alfa, C. E., Hyams, J. S., and Beach, D. H. (1989). The fission yeast cdc2/cdc13/suc1 protein kinase: regulation of catalytic activity and nuclear localization. Cell 58, 485–497.

Chen, Y., Zhao, G., Zahumensky, J., Honey, S., and Futcher, B. (2020). Differential Scaling of Gene Expression with Cell Size May Explain Size Control in Budding Yeast. Mol Cell 78, 359-370.e6.

Chethan, S. G., Rogers, J. M., Vijayakumari, D. et al. (2025). A distinct phase of cyclin B (Cdc13) nuclear export at mitotic entry in Schizosaccharomyces pombe. Open Biol 15, 250199.

Creanor, J., and Mitchison, J. M. (1996). The kinetics of the B cyclin p56cdc13 and the phosphatase p80cdc25 during the cell cycle of the fission yeast Schizosaccharomyces pombe. J Cell Sci 109, 1647–1653.

Curran, S., Dey, G., Rees, P., and Nurse, P. (2022). A quantitative and spatial analysis of cell cycle regulators during the fission yeast cycle. Proc Natl Acad Sci U S A 119, e2206172119.

D’Ario, M., Tavares, R., Schiessl, K. et al. (2021). Cell size controlled in plants using DNA content as an internal scale. Science 372, 1176–1181.

Dischinger, S., Krapp, A., Xie, L., Paulson, J. R., and Simanis, V. (2008). Chemical genetic analysis of the regulatory role of Cdc2p in the S. pombe septation initiation network. J Cell Sci 121, 843–853.

Dorsey, S., Tollis, S., Cheng, J. et al. (2018). G1/S Transcription Factor Copy Number Is a Growth-Dependent Determinant of Cell Cycle Commitment in Yeast. Cell Syst 6, 539-554.e11.

Facchetti, G., Knapp, B., Flor-Parra, I., Chang, F., and Howard, M. (2019). Reprogramming Cdr2-Dependent Geometry-Based Cell Size Control in Fission Yeast. Curr Biol 29, 350-358.e4.

Fantes, P. A., Grant, W. D., Pritchard, R. H., Sudbery, P. E., and Wheals, A. E. (1975). The regulation of cell size and the control of mitosis. J Theor Biol 50, 213–244.

Forsburg, S. L., and Rhind, N. (2006). Basic methods for fission yeast. Yeast 23, 173–183.

Herring, A. J. A Study of Induced Delay in the Division of the Yeast Schizosaccharomyces pombe. University of Edinburgh; 1974. p. Dissertation.

Houser, J. R., Ford, E., Chatterjea, S. M. et al. (2012). An improved short-lived fluorescent protein transcriptional reporter for Saccharomyces cerevisiae. Yeast 29, 519–530.

Keifenheim, D., Sun, X. M., D’Souza, E. et al. (2017). Size-Dependent Expression of the Mitotic Activator Cdc25 Suggests a Mechanism of Size Control in Fission Yeast. Curr Biol 27, 1491-1497.e4.

Lianga, N., Williams, E. C., Kennedy, E. K. et al. (2013). A Wee1 checkpoint inhibits anaphase onset. J Cell Biol 201, 843–862.

Liu, D., Vargas-García, C. A., Singh, A., and Umen, J. (2023). A cell-based model for size control in the multiple fission alga Chlamydomonas reinhardtii. Curr Biol 33, 5215-5224.e5.

Marguerat, S., Schmidt, A., Codlin, S. et al. (2012). Quantitative analysis of fission yeast transcriptomes and proteomes in proliferating and quiescent cells. Cell 151, 671–683.

Miller, K. E., Vargas-Garcia, C., Singh, A., and Moseley, J. B. (2022). The fission yeast cell size control system integrates pathways measuring cell surface area, volume, and time. bioRxiv

Moreno, S., Nurse, P., and Russell, P. (1990). Regulation of mitosis by cyclic accumulation of p80cdc25 mitotic inducer in fission yeast. Nature 344, 549–552.

Mueller, F., Senecal, A., Tantale, K. et al. (2013). FISH-quant: automatic counting of transcripts in 3D FISH images. Nat Methods 10, 277–278.

Murray, A. W., Solomon, M. J., and Kirschner, M. W. (1989). The role of cyclin synthesis and degradation in the control of maturation promoting factor activity. Nature 339, 280–286.

Ohira, M., and Rhind, N. (2022). pomBseen: An Automated Pipeline for Analysis of Fission Yeast Images. bioRxiv

Ohira, M. J., Hendrickson, D. G., McIsaac, R. S., and Rhind, N. (2017). An estradiol-inducible promoter enables fast, graduated control of gene expression in fission yeast. Yeast 34, 323–334.

Padovan-Merhar, O., Nair, G. P., Biaesch, A. G. et al. (2015). Single mammalian cells compensate for differences in cellular volume and DNA copy number through independent global transcriptional mechanisms. Mol Cell 58, 339–352.

Pan, K. Z., Saunders, T. E., Flor-Parra, I., Howard, M., and Chang, F. (2014). Cortical regulation of cell size by a sizer cdr2p. Elife 3, e02040.

Patterson, J. O., Rees, P., and Nurse, P. (2019). Noisy Cell-Size-Correlated Expression of Cyclin B Drives Probabilistic Cell-Size Homeostasis in Fission Yeast. Curr Biol 29, 1379-1386.e4.

Prescott, D. M. (1956). Relation between cell growth and cell division. II. The effect of cell size on cell growth rate and generation time in Amoeba proteus. Exp Cell Res 11, 86–94.

Rhind, N. (2018). Cell Size Control via an Unstable Accumulating Activator and the Phenomenon of Excess Mitotic Delay. Bioessays 40, 1700184.

Rhind, N. (2021). Cell-size control. Curr Biol 31, R1414–R1420.

Russell, P., and Nurse, P. (1987). Negative regulation of mitosis by wee1+, a gene encoding a protein kinase homolog. Cell 49, 559–567.

Schmoller, K. M., and Skotheim, J. M. (2015). The Biosynthetic Basis of Cell Size Control. Trends Cell Biol 25, 793–802.

Schmoller, K. M., Turner, J. J., Koivomagi, M., and Skotheim, J. M. (2015). Dilution of the cell cycle inhibitor Whi5 controls budding-yeast cell size. Nature 526, 268–272.

Schneider, C. A., Rasband, W. S., and Eliceiri, K. W. (2012). NIH Image to ImageJ: 25 years of image analysis. Nat Methods 9, 671–675.

Schulz, H., Flexeder, C., Behr, J. et al. (2013). Reference values of impulse oscillometric lung function indices in adults of advanced age. PLoS One 8, e63366.

Sigrist, S., Jacobs, H., Stratmann, R., and Lehner, C. F. (1995). Exit from mitosis is regulated by Drosophila fizzy and the sequential destruction of cyclins A, B and B3. EMBO J 14, 4827–4838.

Sun, X. M., Bowman, A., Priestman, M. et al. (2020). Size-Dependent Increase in RNA Polymerase II Initiation Rates Mediates Gene Expression Scaling with Cell Size. Curr Biol 30, 1217-1230.e7.

Surana, U., Amon, A., Dowzer, C. et al. (1993). Destruction of the CDC28/CLB mitotic kinase is not required for the metaphase to anaphase transition in budding yeast. EMBO J 12, 1969–1978.

Swaffer, M. P., Kim, J., Chandler-Brown, D. et al. (2021). Transcriptional and chromatin-based partitioning mechanisms uncouple protein scaling from cell size. Mol Cell 81, 4861-4875.e7.

Thormar, H. (1959). Delayed division in tetrahymena pyriformis induced by temperature changes. C R Trav Lab Carlsberg 31, 207–225.

Torres-Garcia, S., Di Pompeo, L., Eivers, L. et al. (2020). SpEDIT: A fast and efficient CRISPR/Cas9 method for fission yeast. Wellcome Open Research 5, 274.

Trcek, T., Chao, J. A., Larson, D. R. et al. (2012). Single-mRNA counting using fluorescent in situ hybridization in budding yeast. Nat Protoc 7, 408–419.

Wang, H., Carey, L. B., Cai, Y., Wijnen, H., and Futcher, B. (2009). Recruitment of Cln3 cyclin to promoters controls cell cycle entry via histone deacetylase and other targets. PLoS Biol 7, e1000189.

Yamano, H., Gannon, J., and Hunt, T. (1996). The role of proteolysis in cell cycle progression in Schizosaccharomyces pombe. EMBO J 15, 5268–5279.

Zatulovskiy, E., Zhang, S., Berenson, D. F., Topacio, B. R., and Skotheim, J. M. (2020). Cell growth dilutes the cell cycle inhibitor Rb to trigger cell division. Science 369, 466–471.

Zhurinsky, J., Leonhard, K., Watt, S. et al. (2010). A Coordinated Global Control over Cellular Transcription. Curr Biol 20, 2010–2015.

